# Inflammation induced Vascular Smooth Muscle Cell expressing Angiopoietin-like 2 to promote the development of abdominal aortic aneurysm

**DOI:** 10.1101/2022.04.12.488011

**Authors:** Guangcan Yang, Jie Guo, Shuang Wu, Hongyou Yu

## Abstract

**Objective:** The objective of this study is to investigate the expression of angiopoietin-like protein 2 (Angptl2) in abdominal aortic aneurysm (AAA) and the expression of Angptl2 in vascular smooth muscle cells (VSMC) under inflammation stimuli.

**Methods:** The mouse AAA model was induced by applying calcium chloride method based the literature and the expression of Angptl2 in AAA was investigated by immunohistochemical staining. The primary VSMCs were cultured and identified following with inflammatory stimuli by Escherichia coli lipopolysaccharide (LPS) for 30 minutes and 24 hours. The expression of Angptl2 were assessed by RT-qPCR and ELISA.

**Results:** The mouse AAA model was successfully established after being induced by calcium chloride. The inflammatory cell infiltration in the arterial tissue of the mice in the experimental group was obvious, and the elastic fiber layer was thinned and broken. Immunohistochemical staining showed that the expression of Angptl2 in AAA was up-regulated; After LPS treatment, both the mRNA and protein expressions of Angptl2 were significantly up-regulated in VSMCs in the experimental group compared with control group.

**Conclusion:** Angptl2 expressed in VSMCs promotes the development of calcium chloride induced AAA.

## introduction

Abdominal aortic aneurysm (AAA) is a pathological dilatation, usually asymptomatic, but very easy to get ruptured [1]. After abdominal aortic rupture, patients may die due to massive blood loss, and the mortality rate is as high as 60%, which is the 13th leading cause of death from the diseases in the United States, and is currently only managed through clinical surgical intervention. Despite the growing number of studies on abdominal aortic aneurysms, there is still lack of the pathogenic explanation for them.

Angiopoietin like 2 (Angptl2) protein is a member of the angiopoietin-like protein family. Angiopoietin-like protein is widely distributed in tissues and organs, including cardiovascular system, lung and kidney [2]. As a member of the angiopoietin family, Angptl2 can promote angiogenesis[3], and Angptl2 secreted by perivascular tissue can promote intimal neogenesis after vascular injury[4]; studies have shown that Angptl2 can promote the polarization of M2 macrophages that is related to inflammation [5], Angptl2 also could maintain tissue homeostasis by promoting adaptive inflammation and subsequent tissue remodeling, while chronic stress-induced overactivation of Angptl2 promotes tissue homeostasis disruption due to chronic inflammation and irreversible tissue remodeling, thereby promoting the development of various metabolic diseases [6]. At the same time, it has been reported that Angptl2 derived from macrophages can promote the development of AAA [7].

Vascular smooth muscle cells (VSMCs) are an important component of the arterial media [8]. The loss of VSMC contractile function is associated with the development of AAA. VSMCs express proteolytic enzymes to degrade the aortic extracellular matrix. Apoptosis of VMSCs is another important factor in the development of AAA [9]. Studies have shown that knockdown of Angptl2 can inhibit the proliferation, migration and invasion of VSMCs [10]. In this study, we found that the up-regulated expression of Angptl2 during the development of AAA and the expression of Angptl2 in VSMC is up-regulated under inflammation.

## Material and Methods

### Materials

The protocol was performed following the guidelines approved by the Institutional Animal Care and Use Committee of Dalian University. Six-moth old C57BL/6 mice and Wistar rats were purchased from Liaoning Changsheng Biological Co., Ltd.; DMED medium was purchased from Wuhan Proceeds; fetal bovine serum was purchased from Gibco; RT-qPCR related reagents were purchased from Hunan Aikerui Biological Engineering Co., Ltd.; Penicillin and trypsin were purchased from Solarbio China; The antibody reagents were purchased from Wuhan Proteitech company in China; primers were designed by Nanjing Jinruisi company;

#### Calcium Chloride induced abdominal aortic aneurysm mouse model

In this experiment, the calcium chloride infiltration method was used to induce mouse abdominal aortic aneurysm model [11]. Briefly, 6-month old male mice were anesthetized with sodium pentobarbital (40mg/kg) then the infrarenal aorta between the left renal vein and aortic bifurcation were treated with CaCl2 (.5M) for 10 mins. The mice were sacrificed on the 42nd and 56th days and the abdominal aorta was dissected, photographed, and store at 4% paraformadehydrate for sectioning and histological staining.

#### Hematoxylin-eosin staining and elastic fiber staining

The mouse arterial tissue was paraffin-embedded for sectioning into hematoxylin-eosin staining (HE staining) and elastic fiber (Verhoeff’s Van Gieson, EVG) staining to detect inflammatory cell infiltration and elastic fiber lesions detection.

#### Immunohistochemical staining

For Angptl2 expression in the aortic wall, the immnunohistalchemistry staining were used to assess the Angptl2 expression. Briefly, the paraffin-embedded sections were dewaxed and rehydrated with serial alcohol concentration, then 5% BSA were used to blocked the section for 1 hour at room temperature followed with membrane penetration with 0.01% Trixton X-100 for 15 mins. The the sections were washed with PBST for 5 mins each for three times. Then primary rabbit anti-Angtplt2 were applied to the section with a dilution 1:1000 at 4°C overnight. Then the sections were washed and applied with Goat anti-rabbit HRP conjugated secondary antibody with a dilution 1:3000 incubating at room temperature for 1 hour. Finally, the DAP was used to bluish the section and the sections were photographed with a Nikon microscope for analysis. The number of Angptl2 cell number were analyzed with ImageJ software.

#### Isolation and culture of smooth muscle cells

Primary vascular smooth muscle cells were cultured by dissecting the rat thoracic aorta and then mincing and digesting the tissue with trypsin EDTA and 200U Collagenase V for 3 hours at 37°C [12]. Then the aortic tissue were applied on the bottom of the tissue cultre flask and cultured in the incubator for 3 days when the VMSC cells will craw out of the tissue. After the smooth muscle cells were cultured and expanded, the passage 5-6 were used to identified by immunofluorescence and used in the following cell experiments

#### Immunofluorescent staining

The VMSC cells were fixed with 100% Methonal at −20°C for 15 mins followed with PBS washing for 3 times. Briefly, the fixed cells were blocked with 10% goat serum for 1 hour at room temperature, then rabbit anti-CNN1 or rabbit anti-Vin were applied with a dilution 1:500 as recommended by the datasheet of the products for 1 hour at room temperature, then the samples were washed with PBST for 3 times, 5 mins each time. The the secondary antibody 1:5000 goat anti-rabbit Coralight 488 was applied for 1 hour at room temperature in dark. DAPI was used to counterstaining the nuclear (Solarbio) for 5min. Images (1600 × 1200 pixels) were acquired at 40x magnification at standardized settings using an Olympus microscope.

#### Real time Quantitive PCR

The smooth muscle cells (2×105) were seeded in petri dishes and cultured for two days, and the serum-free medium was replaced 6 hours before LPS induction treatment, and then returned to normal medium. The cells were treated with LPS for half an hour and the mRNA were collected and used for RT-qPCR assessing the expressing of Angptl2 mRNA. Briefly, the cells total RNA were extracted using the TRIzol reagent according to the manufacturer. Reverse transcriptions were performed by using high-capacity cDNA reverse transcription kit. PCR was performed using the SyberGreen system according to the manufacturer. The following primers used for real-time-PCR: Angptl2 forward: AAGAGCACTGCCAACGTGTA and reverse: CTCTCCATGGACCTGATGGC; GAPDH forward: CTCTCTGCTCCTCCCTGTTC and reverse: CGATACGGCCAAATCCGTTC. mRNAs levels were normalized to glyceraldehyde 3-phosphate dehydrogenase (GAPDH). Changes in expression were calculated using the ΔΔCt method.

#### ELISA

The cells were treated with LPS for 24 hours and then the supernatant were collected for ELISA assessing the Angptl2 expression. Briefly, the supernatant were centrifuged and then 100uL was diluted with equal amount of carbonate buffer and then were coated in the wells of PCT microtiter plate at 4°C overnight. Then the well were washed with PBS for three times followed with a block with 5% goat serum at least 2 hours. Then 100uL of 1:1000 Angptl2 primary antibody were added to each well and incubated for at least 2 hours at room temperature followed with a wash of the plate four time with PBS. The TMB system is used for the detection. TMB solution was added to each well and incubated for 15 min then equal volume of stopping solution was added and the optical density were detected at 450nm

### Statistical Analysis

The statistical analysis were performed with the free statistical software GNU PSPP. Two-sample *t*-test was used to analysis the statistical difference. Data are presented as mean ± SD and P<0.05 was considered statistically significant.

## Results

### The Calcium Chloride induced AAA

Abdominal aortic aneurysm was successfully induced by calcium chloride infiltration method Mice were euthanized on the 42nd day and 56th day after induction of CaCI2, and the diameter of the blood vessels was measured and statistically analyzed after the abdominal aorta was completely dissected (Fig.1). The results showed that the abdominal aorta was significantly enlarged (P<0.001) after induction with CaCI2 for 42 days (n=12) and 56 days (n=12) compared with the control group

**Fig.1.**
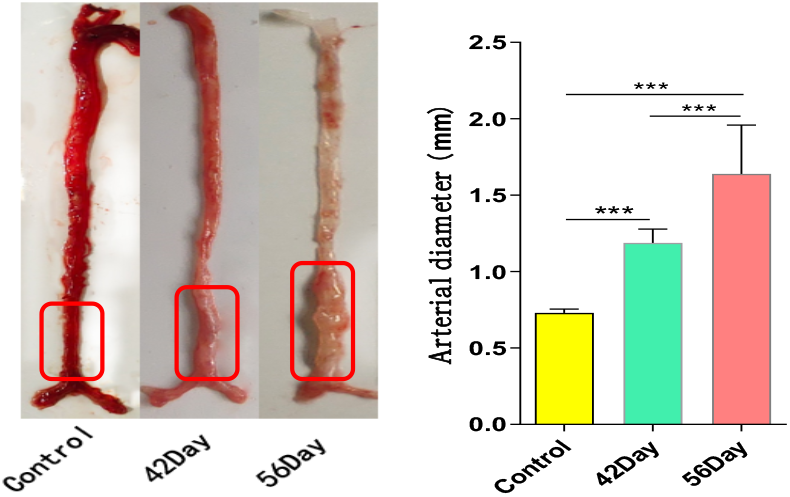
The dissected whole aorta and the aortic diagram analysis. A. The represented whole aorta from control, 46 and 56 days of the calcium chloride induced AAA; B. Assessing the maxium aortic diameter showed that the aortic diameter has been significantly dilated over time courses. ***P<0.0005.

### EVG and HE staining reveals the characters of the aortic wall

The arterial tissue was stained with EVG and HE and photographed under a microscope (Figure 2). The results showed that the elastic fibers in normal arterial tissue were arranged in an orderly manner, and there was no obvious inflammatory cell infiltration; at 42 days after the calcium chloride induction, the elastic fibers of the arteries were thin and irregularly arranged, and the inflammatory cells were significantly infiltrated in the aortic wall. the elastic fiber layer was significantly broken in the arterial group after induction treatment with calcium chloride for 56 days, and the elastic fibers were thin and irregular, and the infiltration of inflammatory cells increased in the arterial group.

**Fig.2.**
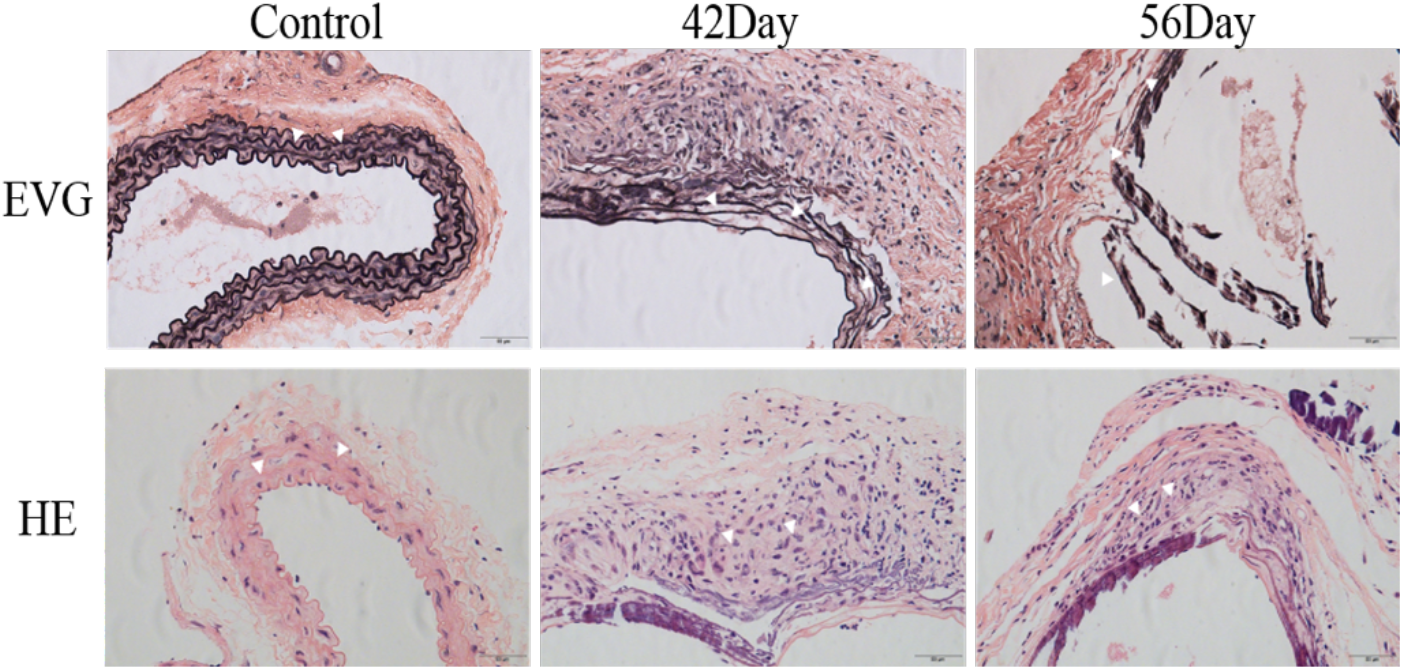
The EVG staining and HE staining of the aortic wall. The figure shows that the elastic fibers were disturbed after 42 and 56 days of calcium chloride induction. The HE staining shows significantly inflammatory cell infiltration in the aortic wall. the magnification 20x and the scale bar length is 1mm

### Increased Angptl2 expression in aneurysms

The aneurysm tissue sections were immunohistochemically stained and photographed under a microscope. It was found that the normal tissue was intact in the arterial group, and there were fewer Angptl2 positive cells; after calcium chloride was administered to induce the arterial tissue structural lesions, the Angptl2 positive cells in the arterial outer layer group increased significantly(Figure 3A). And on the 56th day, we noticed that there were relatively few Angptl2-positive cells in the relatively intact inner layer of the artery, while the number of Angptl2-positive cells increased significantly in the peripheral tissue where the inner layer structure was destroyed (Figure 3B).

**Fig.3.**
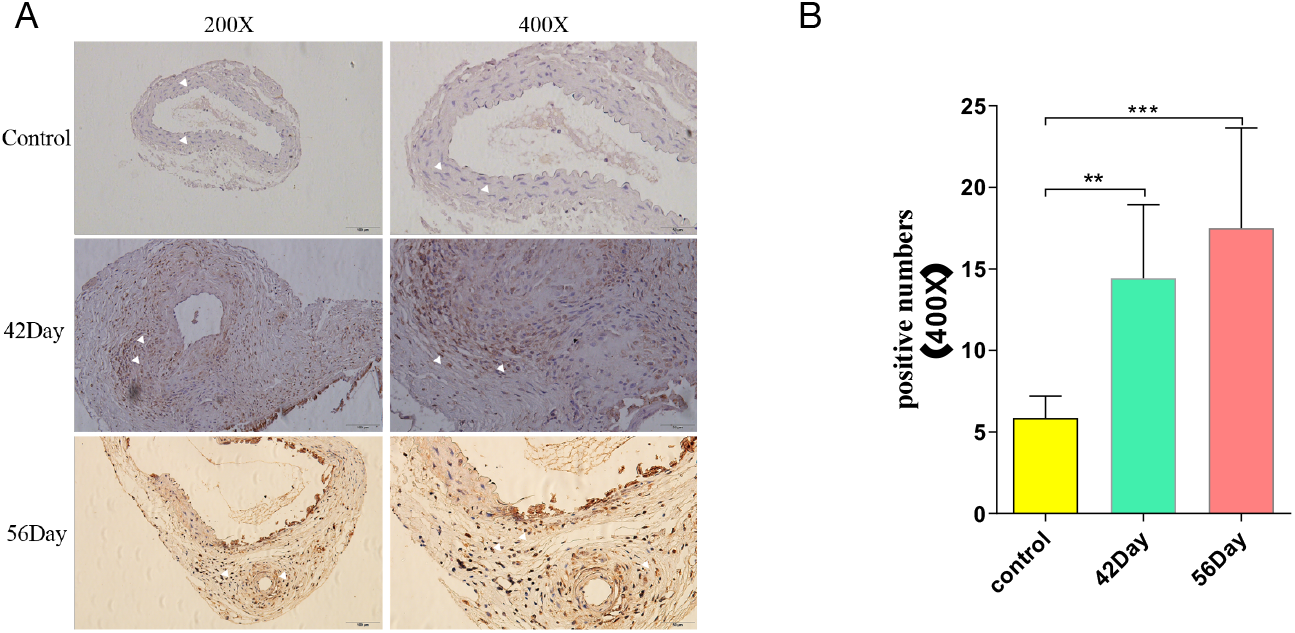
The Expression of Angptl2 in the aortic wall. A. The immunohistochemically stained aortic section in control, 42 and 56 days of calcium chloride induction and showed significant Angtpl2 positive cells in the aortic wall (the white arrow); B. By calculating the positive numbers of Angptl2 cells in the aortic wall showed that there is significantly higher number of Angptl2 positive cells both in the 42 days and 56 days group. the magnification 20x and the scale bar length is 1mm **P<0.005, ***P<0.0005

### Increased Angptl2 mRNA and protein expression in VSMCs

As the VMSCs are the major cells in the aortic wall, so we cultured primary VMSCs to assess whether the inflammation induces the expression Angptl2 in the VMSCs. The VMSCs were culture using rat aortic tissue and the cells were fully grew in 5 days. The immunofluorescent staining shows that the cells were expressing the VMSC specific marker both CNN and Vin, which indicates that the cultured cells were VMSCs(Figure 4)

**Fig.4.**
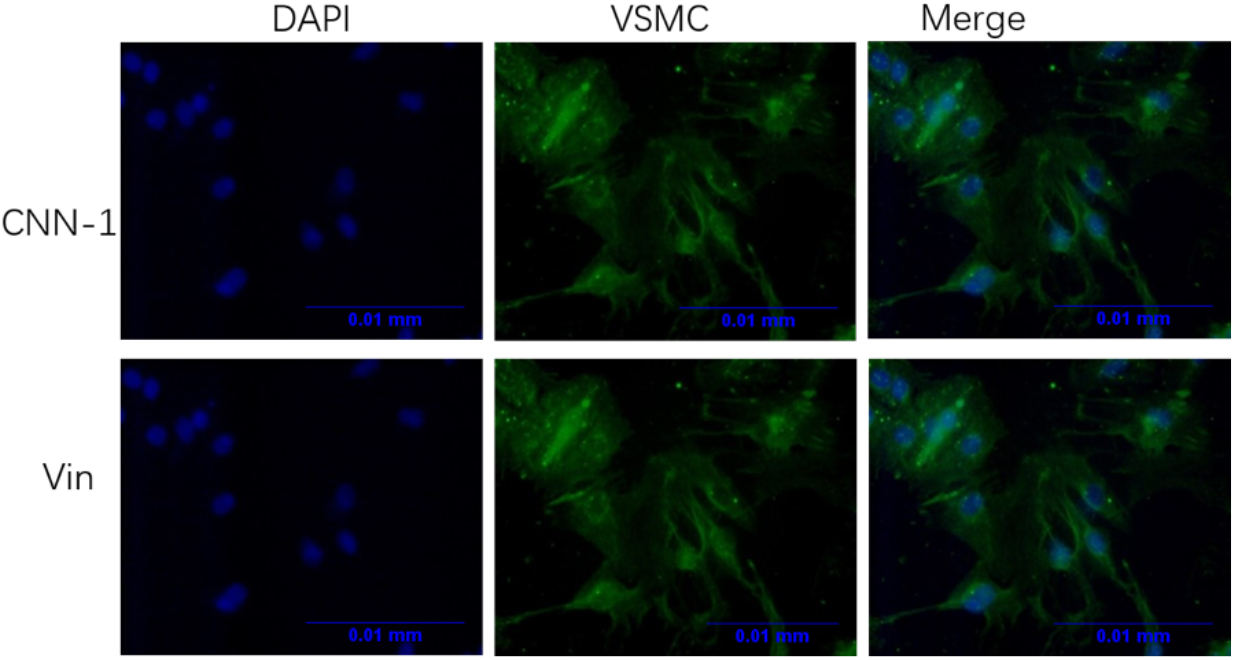
Identification of VSMCs by immunofluorescence staining. The immunofluorescent staining reveals that the primary culture cells expressed both the CNN1 and Vin which indicates that the culture cells were purified VMSCs. magnification 20x and the scale bar length is 0.01mm.

Then the VSMCs were stimulated with Escherichia coli lipopolysaccharide (LPS) for 30 minutes and 24 hours to extract RNA and to collect supernatant respectively for RT-qPCR and EILSA for assessing the expressing of Angptl2. The results showed that the mRNA expression of Angptl-2 in VSMC was significantly up-regulated after 30 minutes of LPS stimulation (Figure 5A). The Angptl2 protein in the supernatant of the cell culture medium was assess by ELISA and it was found that the LPS induced significantly Angptl2 secretion compared with the control group (Figure 5B).

**Fig.5.**
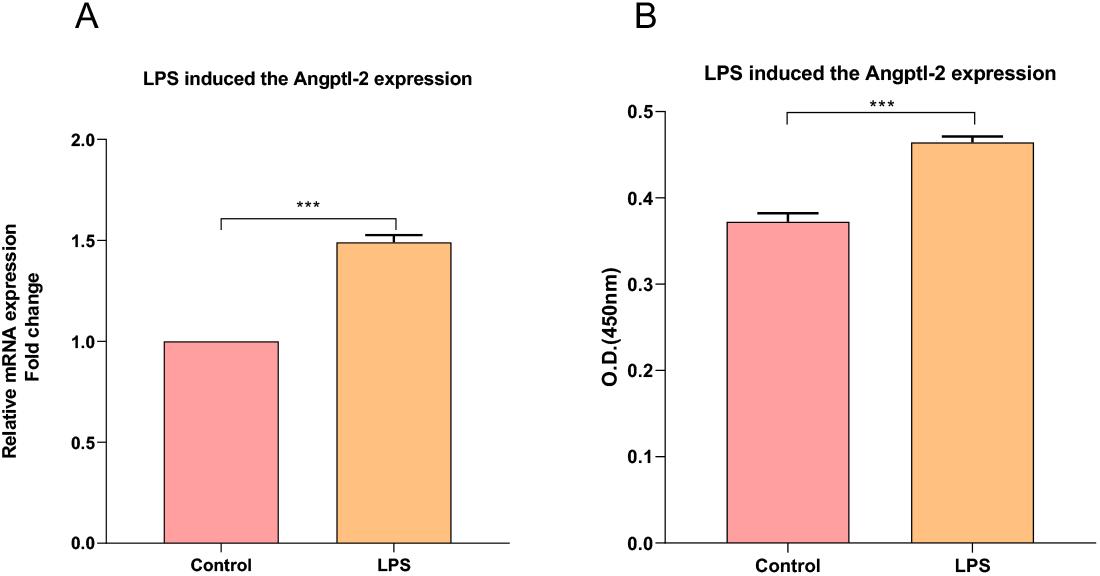
LPS-induced Angptl2 expression in VSMCs. A. The mRNA expression in the VMSCs were significantly up-regulated induced by the LPS compared with the comtrol group. B. The secreted Angptl2 were significantly higher in the LPS inducted group compare the control group. ***P<0.0005

## Discussion

The drug treatment and prevention of abdominal aortic aneurysm disease is still a difficult problem to be solved. Surgical treatment is only suitable for some aneurysms. For small aneurysms or other patients who are not suitable for surgery, only conservative treatment can be selected, but there are still as many as 70%. of small aneurysms continue to expand to rupture and are expected to be related to multiple factors after surgery [13, 14]. In the development of abdominal aortic aneurysm, VSMC phenotype function conversion and dysfunction, the VSMC phenotype can be converted into macrophage-like cells during the development of abdominal aortic aneurysm, and the phenotype conversion is correlated with arterial calcification and arteriosclerosis density [9].

In this study, a mouse AAA model was constructed by CaCl2 induction, and the study showed that Angptl2 expression was increased in AAA tissue. Angptl2 is involved in the regulation of inflammatory responses. Studies have shown that knockdown of Angptl2 inhibits LPS-induced acute inflammation in the eye [15], and Angptl2 regulates the inflammatory response of human gingival epithelial cells [16]. The expression of Angptl2 was increased, and the experimental results indicated that the upregulation of Angptl2 in VSMC was related to the development of AAA. Studies have shown that macrophage-derived Angptl2 can promote the development of AAA, and the loss of Angptl2 inhibits the development of AAA induced by CaCI2 [7]; Angptl2 is associated with chronic inflammation and atherosclerosis [17], Angptl2 can induce macrophage and Leukocytes adhere to the vessel wall [18] and can also accelerate atherosclerotic calcification [19]. Downregulation of Angptl2 reduces monocyte recruitment and inflammation in the endothelium, and also stimulates endothelial repair to alleviate atherosclerosis [20]. AAA is associated with atherosclerosis [21], and the aorta in patients with AAA is more inflamed than in patients with atherosclerosis, and the calcification within the aneurysm is more pronounced [22]. We speculate that Angptl2 has a similar role in AAA development, and that Angptl2 in perivascular adipose tissue has a pathological role in vascular remodeling [23].

Angptl2 may promote angiogenesis during the development of AAA. Angptl2 is an orphan ligand [24, 25] that inhibits endothelial barrier leakage [24]. Studies have shown that Angptl2 can promote endothelial cell sprouting and angiogenesis by regulating endothelial colony cell differentiation [3]. Angiogenesis alters the vascular barrier and permeability, and the formation of new blood vessels can mediate the entry of inflammatory factors and increased infiltration of macrophages [26].

Matrix metalloproteinases (MMPs) are a class of zinc-dependent endopeptidase proteins that can degrade the extracellular matrix and promote the development of AAA. Studies have shown that MMP-1, MMP2, and MMP-9 are up-regulated during the development of AAA [27, 28]. MMP-2 is mainly derived from smooth muscle cells and fibroblasts, and a small part is derived from macrophages, while MMP-9 is mainly derived from macrophages and a small part is derived from neutrophils [29-31].

Studies have shown that Angptl2 increases the expression of pro-inflammatory cytokines and MMP-9 by activating an NF-κB-dependent cascade in infiltrating macrophages, thereby accelerating chronic inflammation and promoting destructive remodeling of the vessel wall leading to AAA development [7]]. Silencing Angpt2 reduces the expression of MMP-9 [32], while overexpression of Angptl2 promotes the expression of MMP-2 and MMP-9 in human osteosarcoma cells (U2OS). Therefore, Angptl2 expressed in VSMC has an important role in the development of AAA.

In summary, Angptl2 expressed by VSMC could participate in the development of AAA through multiple pathways. Angptl2 plays a very important role in the development of AAA, and it also suggests that Angptl2 can be used as a potential therapeutic target for the development of AAA drugs.

## Conclusion

Angptl2 plays critical roles in the development of AAA and the expression Angptl2 in the VMSCs under inflammatory stimuli could contribute the dilation of the aorta.

